# Variability in fitness effects and the limitations of fitness optimization

**DOI:** 10.1101/107847

**Authors:** Christopher J Graves, Daniel M Weinreich

**Affiliations:** Brown University, Department of Ecology and Evolutionary Biology and Center for Computational and Molecular Biology. Providence, RI, USA

**Keywords:** Natural selection, Fitness optimization, Varying environments, Group selection, Inclusive fitness, Mutation rate

## Abstract

Evolutionary biologists commonly assess the evolutionary advantage of an allele based on its effects on the lifetime survival and reproduction of individuals. However, alleles affecting traits like sex, evolvability, and cooperation can cause fitness effects that depend heavily on differences in the environmental, genetic, and social context of individuals carrying the allele. This variability makes it difficult to summarize the evolutionary fate of an allele based solely on its effects on any one individual. In this review we show how attempts to average over variability in the fitness effects of an allele can sometimes cause misleading results. We then describe a number of intriguing new evolutionary phenomena that have emerged in studies that explicitly model the fate of alleles that influence long-term lineage dynamics. We conclude with prospects for generalizations of population genetics theory and discuss how this theory might be applied to the evolution of infectious diseases.

## I. Introduction

Evolution by natural selection is driven by heritable differences in the reproductive success of individuals. However, the long-term outcome of natural selection depends not only on the effects of an allele on individual bearers but also on its effects across its entire lineage of descendants-defined here as the genealogy of an allele from its origination to its ultimate fixation or extinction in the population (Sidebar 1). When fitness effects are invariant across a lineage, the long-term fate of an allele can be deduced in a relatively straightforward manner from its recursive effects on survival and reproduction across descendent members of the lineage. In other cases, the evolutionary success of an allele is not an obvious consequence of its effects on individuals. For example, variable environments can cause the same allele to have differing effects on fitness depending on an individuals’ environmental context. Similarly, fitness effects may vary due to the presence of other alleles in the genome, which are themselves polymorphic in the population. In such cases, it is often presumed that traits will tend to spread by natural selection so long as they are beneficial to their carriers on average (Eshel 1973, Nunney 1999). This implies that natural selection favors traits that are beneficial not strictly to individuals, but to genetic lineages as a whole.

The concept that natural selection may optimize quantities related to the average success of an allele across a lineage has arisen in a wide range of problems ranging from varying environments to the evolution of sex and cooperation (Akçay and Van Cleve 2016, Eshel 1973, Kussell and Leibler 2005, Lehmann, et al. 2016, Nunney 1999, Nunney 1999). In general, this idea arises when the fitness effect of an allele varies between individual carriers, thereby limiting the ability to infer the long-term success of an allele based on measures of individual fitness alone. A large class of evolutionary problems fit this description and they can be classified by whether the variability across a lineage arises due to environmental, genetic, or social factors. We outline examples of each in Table 1 and describe them in more detail in the main text. Each source of variation has largely been discussed within its own body of literature, where equivalent concepts are used to describe a distinct set of adaptations, often with distinct terminology. Despite some obvious similarities, there have been few attempts at synthesizing what is known in each of these cases into a formal quantitative theory of the evolution of alleles with lineage-variable fitness effects.

**Table 1.** Sources of variability across a lineage and associated adaptations (*Typeset near* ‘Introduction’)

|  | Basis of variability in fitness |  |  |
| --- | --- | --- | --- |
|  | Environmental | Genetic | Social |
| Specific examples | Spatial variation | Sex | Kin selection |
|  | Temporal variation | Mutation | Multi-level selection |
|  |  | Clonal interference |  |
|  |  | Epistasis |  |
| Adaptations | Bet-hedging | Sex/recombination | Cooperation |
|  | Phenotypic plasticity | Mutation rate modifiers | Policing |
|  |  | Evolvability |  |

Averaging the fitness effects of an allele across a lineage shifts the target of adaptation from individuals to lineages. However, one must acknowledge possible limitations in the ability of natural selection to favor traits that confer a long-term benefit to a lineage. Specifically, natural selection is myopic in nature- acting to increase the frequency of traits that confer an immediate advantage to individuals without regard to their future utility to a lineage. This shortsightedness can have dramatic consequences, particularly if it results in the permanent extinction of an allele prior to it realizing any average benefit in the long-term. Indeed, the notion that natural selection will act most strongly on alleles that confer a short-term advantage was championed by Maynard Smith (1964) and Williams (1966) in their now famous critique of group selection, and is still in use (Lynch 2007, Sniegowski and Murphy 2006). When does natural selection favor traits that confer a long-term benefit averaged across a lineage and when does shortsighted selection limit this ability?

After briefly summarizing results from classical, lineage-invariant theory that successfully relates individual fitness to a lineage’s eventual fate, we discuss a diversity of examples of lineage-variable fitness, i.e., cases in which the fitness effects of an allele vary across its lineage of descendants. We illustrate the shortcomings of averaging variability across the lineage in finite populations, in which alleles that are beneficial in the long-term are nevertheless vulnerable to extinction. Consequently, shortsighted selection in finite populations can limit the ability of natural selection to optimize even these measures of fitness. Finally, we discuss other counterintuitive results that emerge in examples where lineage-variable fitness is modeled explicitly. These results show that the fate of an allele can be sensitive not only to its fitness effects across a lineage, but to features unrelated to classical notions of fitness, such as the population size. We conclude by highlighting implications for the evolution of infectious diseases and directions for future work.

## II. Lineage-Invariant Fitness Effects

Evolutionary biologists are fundamentally concerned with understanding the outcome of natural selection on traits that influence the survival and reproduction of their carriers. Before discussing cases in which the fitness effects of an allele are variable across its lineage we first consider the case where fitness effects are invariant across a lineage. Our approach throughout will be on the field of population genetics, which has a rich tradition of analyzing dynamical models that combine various evolutionary forces including natural selection, genetic drift, and mutation. Such dynamical treatments of evolution provide a comprehensive analysis of a lineage – starting from its origination in the population and ending with its ultimate fixation or extinction. We will therefore be decidedly brief in our overview of other aspects of evolutionary theory, which include techniques such as game theory and quantitative genetics.

Consider an allele that influences the expected number of surviving offspring produced over the lifetime of its carriers. Formally, we allow the precise number of offspring produced by any particular individual in this lineage to be a Poisson random variable drawn independently from an identical distribution, with mean defined as the Wrightian fitness, ***w***. This concept of fitness articulates well with the Darwinian notion of fitness as lifetime reproductive success. The most fundamental consideration regarding the fate of an allele by natural selection is whether the allele influences this measure of fitness relative to the resident “wild type” in the population. In population genetics, this fitness effect is most often denoted with the selection coefficient, *s*, defined as the proportional change in expected number of offspring relative to the wild-type: *s* ≡ *w*_mut_/*w*_wt_ – 1.

Now consider a population with constant size, *N*. Since the number of surviving offspring born to an individual is a random variable, we allow for random fluctuations in the number of individuals carrying an allele as the basis for genetic drift. Assuming that generations are discrete and non-overlapping and approximating the Poisson offspring distribution with a binomial so that the population size remains fixed, we can describe the allele frequency dynamics using the Wright-Fisher model. We emphasize that the Wright-Fisher process and related models capture the interplay between natural selection and genetic drift in finite populations by incorporating stochasticity in the number of surviving offspring born to each individual. However, by definition all models of lineage-invariant selection assume that the distribution in that number remains constant across the lineage (Figure 1A).

**Figure 1.**
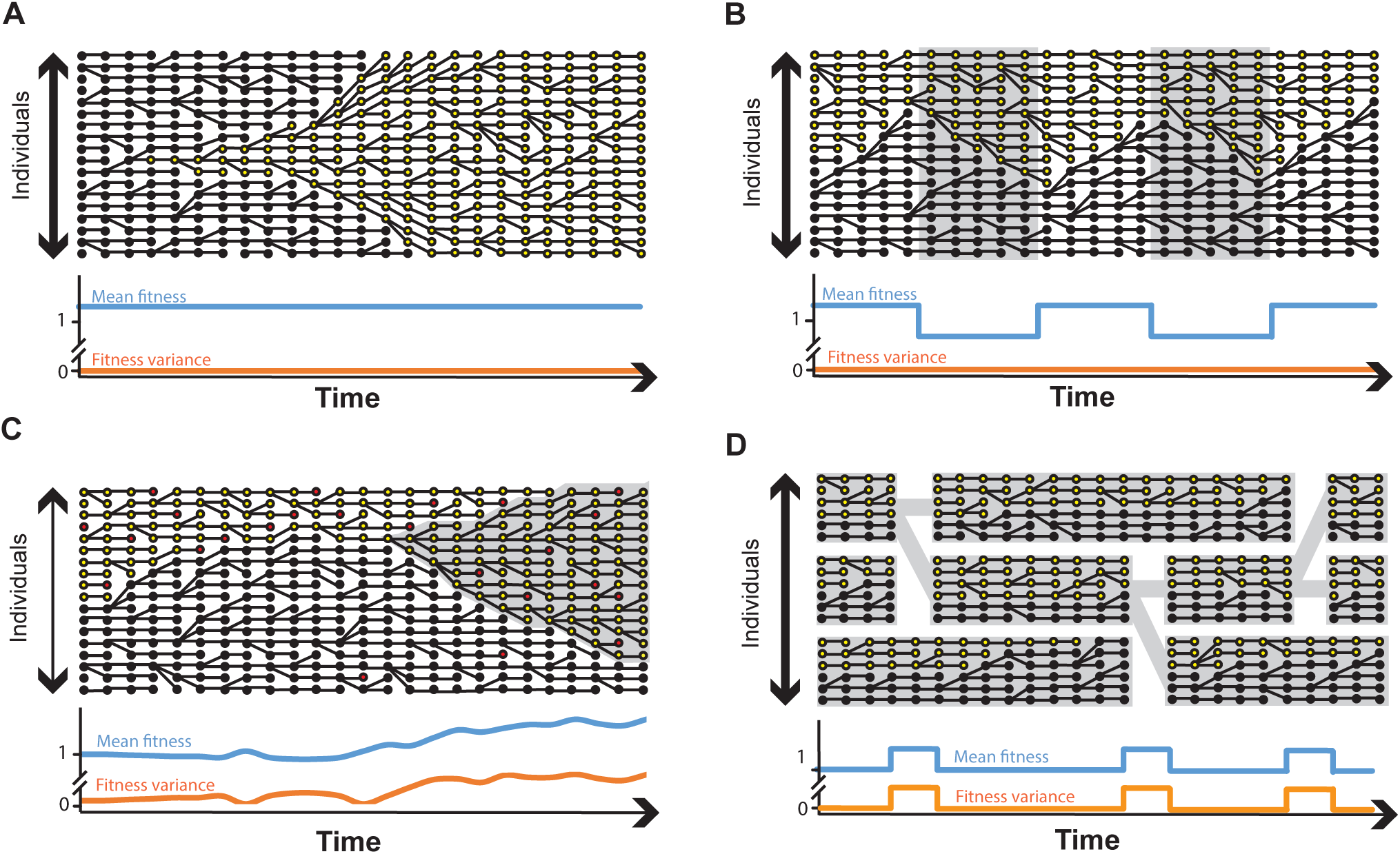
Variability in fitness across a lineage in diverse models. A large number of realistic biological scenarios can result in the presence of variation in fitness across a lineage either among contemporary individuals (vertical axis) or between individuals in time (horizontal axis). Genealogies are shown for two competing allelic lineages indicated by circles. The focal lineage is shaded yellow and the wild-type lineage is shaded black. Curves on the bottom of each panel depict changes to the mean (blue) and variance (orange) of fitness across the focal lineage. **A**. Lineage carrying a beneficial allele (yellow) rising to fixation under the classical scenario of lineage invariance. **B**. Lineage carrying an allele that alternates from beneficial to deleterious in a variable environment. Contemporary individuals share an identical fitness, and hence an identical selection coefficient, but this quantity changes over time. **C**. Evolution of a mutator lineage that experiences increased rates of both deleterious (red dots) and beneficial (grey background) mutations. **D**. A cooperative lineage under a group selection model. Within-group selective pressures cause the allele to be disfavored over short timescales. Groups with more cooperative alleles tend to displace other groups over longer timescales (shown with solid grey lines). In both **C** and **D**, fitness in the lineage varies both among contemporary individuals and over time.

Given this framework, we can obtain solutions for a number of quantities pertaining to the fate of a mutant allele based on its selection coefficient, *s*. Of particular interest given our concern with the ultimate fate of a lineage is the probability that an allele eventually displaces all alternatives in the population, known as the probability of fixation, *P_fix_*. Kimura (Kimura 1962) found this quantity for a mutation starting at frequency *x_0_*, in a haploid, randomly mating population of size *N*, using a continuous diffusion approximation of the Wright-Fisher process:

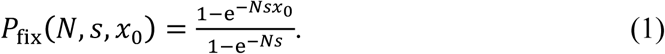
This result highlights many of the key features of classical population genetics theory. Solving for the limit as *s* approaches zero leads to *P_fix_* = *x*_0_. This defines the neutral expectation that the probability of fixation of an allele is simply equal to its starting frequency. Focusing on the case where an allele starts from a single mutation in the population, we take *x_0_* = 1/*N*. Now consider a beneficial mutation, *s* > 0. Here, *P_fix_* > 1/*N*, but only asymptotically approaches *s*, even as population size *N* tends to infinity (Haldane 1927). In other words, fixation of even a strongly beneficial mutation is not assured, reflecting the fact genetic drift dominates allele frequency dynamics until there are roughly 1/*s* copies in the population. This effect is worsened in small populations since 1/*s* copies may be an appreciable fraction of the population. Thus as s or *N* get small, 1/*s* approaches *N* and genetic drift comes to dominate selection. This result implies that mutations are effectively neutral from the standpoint of natural selection, unless *s* > 1/*N*. Finally, and somewhat less intuitively, Kimura’s formula also shows that even deleterious mutations (*s* <0), can have a nonzero fixation probability. Here again, genetic drift can overwhelm natural selection in populations roughly no larger than 1/|*s*| individuals.

## III. Lineage-Variable Fitness Effects

Under the assumption that an allele exerts a constant, lineage-invariant effect on fitness, Equation 1 demonstrates that a mutant’s fitness effect is sufficient to predict the fate of its lineage. We now turn to cases where variability in the fitness effects of an allele can cause this result to fail. Examples of lineage-variable fitness effects emerge under many realistic biological scenarios, where alleles do not act alone to influence fitness but interact with different environmental, genetic, or social factors (Table 1, Figure 1). Consequently, the number of offspring produced by individuals in a lineage may not be drawn from any fixed distribution, violating the assumption of lineage invariance underlying Equation 1. We emphasize that such variability in offspring number is beyond that captured in models like Wright-Fisher, which typically require the distribution of offspring number to be fixed. Our goal in this section is to highlight some of the relevant examples of variability in fitness of an allele represented by the three classes in Table 1, and to build some intuition for how they have been handled in the literature. We also seek to show that adaptations associated with each example depend uniquely on the effects of an allele on the fate of a lineage rather than on individual success.

### Environmental interactions

Natural environments are inherently variable and therefore present an obvious challenge to the assumption that an allele will have the same effect on fitness for all members of a lineage.

Variation in the environment over time will cause contemporary members of a lineage to experience the same distribution of offspring number, but this distribution now depends on time (Figure 1B). Contrastingly, under spatial variation in the environment, contemporary members will experience fitness effects that depend on the interaction between their shared allele and the local environment they encounter. This again implies that no single distribution in offspring number will be generally applicable. In either case, if environmental change is so rapid that individuals encounter a succession of different environments in their lifetime, then fitness can be described as a lifetime average of total survival and reproduction (Levins 1968). We will therefore focus on the more interesting case where environments vary on a timescale greater than the generation time of the organism; here averaging can often be misleading.

The greatest progress has been made in models of temporally varying environments, in which case the selection coefficient *s* is no longer a constant, but a time-dependent quantity, *s*(*t*). Formal analysis typically requires specifying a particular form of *s*(*t*) at the expense of generality. It is commonly assumed that environments are randomly drawn from a fixed distribution or that that the population size is infinite (Dempster 1955, Gillespie 1973, Kussell and Leibler 2005, Lewontin and Cohen 1969). Under these assumptions, a diverse set of models can be integrated based on how variation in fitness correlates within and between members of two competing lineages (Frank and Slatkin 1990). We note, however, that such an approach is limited to deriving the instantaneous change in allele frequency rather than explicitly modeling lineage dynamics. Another consequence of assuming random environmental change and infinite populations is that natural selection will favor alleles that increase the long-term growth rate of a lineage, averaged over all environments (Dempster 1955, Gillespie 1973, Kussell and Leibler 2005, Lewontin and Cohen 1969, Stearns 2000). Formally, this corresponds to an increase in the geometric mean fitness, or equivalently, the mean intrinsic growth rate (Sidebar 2), and is generalizable to other forms of *s*(*t*) (Cvijović, et al. 2015). Importantly, even arbitrarily large but finite populations experience genetic drift, which can limit the ability to maximize the long-term growth rate of lineages in certain environmental scenarios. We discuss these limitations in more detail below.

Despite its limitations in finite populations, the principle that in variable environments natural selection acts to increase geometric mean fitness is a key theoretical insight, and it is presumed to underlie numerous adaptations. These include strategies like developmental and phenotypic plasticity that allow adaptive phenotypic responses to environmental conditions that may not be encountered by all individuals (Meyers and Bull 2002, Via, et al. 1995). Most notable is the evolution of bet-hedging traits in which an allele causes the exaggeration of phenotypic noise among members of a lineage, thereby allowing a single genotype to spread environmental risk among different phenotypes that are suited to different environments (Fraser and Kaern 2009, Gillespie 1974, Kussell and Leibler 2005, Philippi and Seger 1989). Such a strategy is inherently dependent on selection favoring traits that confer a long-term benefit across a lineage, since individuals will experience differing fitness values depending on their phenotype and the environment they encounter. By spreading the risk of fitness losses under future environmental uncertainty across members of a lineage, bet-hedging helps to ensure survival and reproduction across the lineage as a whole, regardless of the environment. Examples of adaptive bet-hedging strategies have been noted in plants (Childs, et al. 2010, Clauss and Venable 2000, Gremer and Venable 2014), insects (Hopper 1999, Menu, et al. 2000), and microbes (Balaban, et al. 2004, Jones and Lennon 2010, Levy, et al. 2012).

### Genetic interactions

Alleles don’t influence fitness alone but do so as part of an integrated genome. The genetic background of an allele is therefore another important source of variability in fitness across a lineage. Perhaps the most obvious example is that of epistasis (Phillips 1998), in which the fitness effect of a mutation depends on its genetic context. Empirical evidence suggests that epistasis among alleles is widespread (Costanzo, et al. 2016, Kryazhimskiy, et al. 2014, Wang, et al. 2014, Weinreich, et al. 2013) and therefore provides an important source of variability in the fitness effects of an allele, particularly in sexual populations. Similar variation in fitness can occur in asexual populations due to secondary mutations that arise on the genetic background of an allele as it spreads. This effect is most important in large populations or under high mutation rates. Such conditions lead to clonal interference (Gerrish and Lenski 1998), in which multiple asexual lineages carry competing beneficial mutations, thereby interfering with one another’s fixation. The fate of a lineage under clonal interference cannot be decided by the selection coefficient of a single allele, but instead depends on the process of successive mutations accumulating along a series of competing asexual lineages (Desai and Fisher 2007, Lang, et al. 2013). Indeed, this presents a major hurdle to evolving asexual populations, since the lack of recombination implies strict genetic linkage among mutations that occur on the same background. This lack of recombination can also lead to Muller’s ratchet (Haigh 1978, Muller 1964), in which the serial fixation of deleterious mutations by genetic drift can cause fitness to erode along an asexual lineage.

The constraints on asexual adaptation due to clonal interference and Muller’s ratchet provide strong arguments for why so many organismal life cycles include periods of recombination or sexual reproduction. These arguments are invariably related to the idea that alleles influencing sex can increase the long-term average evolutionary success of a lineage (Nunney 1999). This is because sex is inherently costly to individuals, who must invest time and energy in mating and further invest resources into the production of males, which are not capable of independent reproduction (Maynard Smith 1978). These costs could, however, be balanced if sex increases the long-term fitness of lineages (Nunney 1989, Nunney 1999). For example, under certain conditions of epistasis, recombination can accelerate both the pace of adaptation (Eshel and Feldman 1970) and the ability of populations to purge deleterious mutations and fend off Muller’s ratchet (Kondrashov 1988). Furthermore, sexual reproduction can increase rates of adaptation by allowing beneficial mutations that arise on different backgrounds to be combined into a single genotype, thereby limiting the constraints imposed by clonal interference (Cooper 2007, McDonald, et al. 2016). Finally, the red-queen hypothesis (Hamilton, et al. 1990, Van Valen 1973), asserts that the constant creation of new genotypes under recombination can allow organisms to more readily compete in a co-evolutionary arms race with parasites. Indeed, sex is likely to have evolved for a combination of reasons and empirical observations support many of the hypotheses that have been put forth (Colegrave 2002, Cooper 2007, Goddard, et al. 2005, McDonald, et al. 2016, Morran, et al. 2011).

Sex and recombination are not the only processes that increase rates of adaptation. There has been substantial recent attention on whether natural selection can act more generally on the ability of populations to adapt, or its evolvability. Selection for evolvability is contentious, since the ability to adapt to future contingencies is a feature of populations and would therefore appear to require evolutionary foresight and group selection operating on biological populations (Lynch 2007, Pigliucci 2008, Sniegowski and Murphy 2006, but see Watson and Szathmáry 2016). However, traits that increase evolvability could also arise by the process of natural selection favoring traits that are beneficial on average, with lineages being more likely to persist over longer evolutionary periods if they are able to adapt to future conditions (Eshel 1973). While numerous traits could increase evolvability (Wagner and Altenberg 1996), there has been a great deal of attention paid to the evolution of alleles which influence the mutation rate – known as mutation rate modifiers (Denamur and Matic 2006, Sniegowski, et al. 2000). Mutation rate modifiers have been observed in microbial populations both in the lab (Sniegowski, et al. 1997) and in nature (LeClerc, et al. 1996, Matic, et al. 1997). The fate of such “mutator” alleles is intriguing, since they often arise without a direct effect on fitness themselves (Chao and Cox 1983, Sniegowski, et al. 1997). In asexual populations, mutators are still physically linked to the mutations they produce and can thereby influence the statistical properties and long-term fate of lineages (Figure 1C). In such scenarios, evolvability arises as a by-product of indirect selection and genetic hitchhiking of mutators (Sniegowski and Murphy 2006). However, there are also notable exceptions in which histories of repeated environmental change could directly favor the evolution of traits that increase evolvability. This appears to be the case in pathogens, where elevated mutation rates in antigens to increase the capacity to adapt to a dynamic vertebrate immune response (Graves, et al. 2013, Moxon, et al. 1994).

### Social interactions

Fitness is influenced not only by environmental and genetic factors but also by interactions with other conspecifics. These interactions can create a type of lineage-variable fitness known as frequency dependent selection, where the fitness effects of an allele are dependent on the frequency of the allele in the population. Frequency dependence is conveniently analyzed in the context of evolutionary game theory (Sidebar 3), which allows one to consider the ability of an initially rare allele to invade a population fixed for a wild-type allele (Maynard Smith 1982, Maynard Smith and Price 1973). This approach provides a generalization of the concept of a selection coefficient to instances where fitness cannot be wholly represented by a constant value. A classic example of frequency-dependent selection arises when considering cooperative traits. Here, cooperative acts incur a cost to individuals and are therefore susceptible to invasion by selfish “cheater” strategies that avoid the cost of cooperating while still reaping the benefit. Cheaters are typically beneficial when rare, since their fitness advantage requires interactions with other cooperators. Despite the inherent susceptibility to cheaters, cooperation is common in nature and is presumed to underlie major transitions in evolutionary history, such as the evolution of multicellularity (Szathmary and Maynard Smith 1995). The mechanisms promoting the evolution and maintenance of cooperation are therefore of long-standing interest to biologists.

Significant theoretical progress on the evolution of cooperation arose with the formulation of inclusive fitness theory. Hamilton (1964) showed that genes controlling cooperation may be beneficial on average so long as the beneficiary of cooperative actions are kin, which are likely to share the genes controlling cooperation by common descent. The key realization of this theory is that cooperative acts need not directly increase the reproductive success of individual bearers, but instead must increase the average effect of a gene across the lineage of cooperators (Akçay and Van Cleve 2016). Cooperation can also be stable under cases of multi-level selection (Luo 2014, Simon, et al. 2013, Traulsen and Nowak 2006). The formation and dissolution of new groups is itself a reproductive process and the long-term fate of a lineage is therefore sensitive to the influence of an allele on group-level reproduction (Figure 1D). A well-known example is infectious diseases, discussed below, in which individual cells or viral particles must replicate within hosts and also spread among hosts to establish new infections.

There is ample empirical evidence for the stability of cooperative traits in nature if selection favors traits that increase the long-term growth rate of a lineage at the expense of individual fitness. For example, a large number of studies have shown how cooperation, which appears costly to individuals, can prevail through the action of group selection and kin selection (Gore, et al. 2009, Koschwanez, et al. 2013, Rainey and Rainey 2003, Turner and Chao 1999, Velicer, et al. 2000). Perhaps more intriguing is the evolution of “policing” phenotypes that function to reduce the short-term benefits of selfish cheater phenotypes and thereby stabilize cooperation (Frank 1995, Nunney 1999, Travisano and Velicer 2004). For example, in social insects, reproduction by the worker caste constitutes a selfish trait that can undermine colony reproductive interests. To prevent selfish reproduction among workers, social insects have evolved anti-cheater strategies, where colony members will systematically destroy eggs laid by workers (Ratnieks and Visscher 1989). Tumor suppressor genes of multi-cellular organisms perform a similar function by recognizing and destroying cells that violate normal growth regulation and thereby preventing outgrowths of genetically selfish cancer cells (Nunney 1999). Finally, group selection dynamics can even result in Simpson’s paradox, in which the overall frequency of cooperators increases despite their systematic tendency to decrease within groups (Chuang, et al. 2009). The fact that a trait can spread even as it selects against in every individual carrier shows the potential for selection on long-term growth rate to prevail over selection on individuals.

## IV. Limitations of fitness averages

### Limitations due to short-sighted selection

A central theme in many of the treatments of lineage-variable fitness effects is that fitness differences can be averaged across a lineage using concepts like geometric mean fitness and inclusive fitness. These extended fitness averages provide a convenient way to determine if an allele has a positive or negative affect on a lineage- by instead determining whether it increases long-term the rate of spread of a lineage as a whole. We also note the equivalency between these concepts and several related averages that depict long-term growth rates. For example, pathogens are widely assumed to maximize their long-term transmission success, *R*_0_ (Alizon, et al. 2009, Anderson and May 1982). Similarly, Lyapunov exponents are sometimes used to derive long-term growth rates in variable environments (Kussell and Leibler 2005) and the concept of invasion fitness in evolutionary game theory (Sidebar 3) indicates whether natural selection tends to favor a trait under frequency dependence (Lehmann, et al. 2016). Similar averages have been used to deal with variation in an allele’s genetic background (Falconer 1994, Livnat and Papadimitriou 2016). In general, averages across the variability in reproductive success are meant to allow one to directly define an “effective” selection coefficient in order to identify which allele increases fitness. An even more ambitious goal would be to salvage Equation 1, as is the case under scenarios of rapid environmental change (Cvijović, et al. 2015).

Unfortunately, there are fundamental problems with the use of these averages that can preclude natural selection from maximizing fitness averages. Specifically, shortsighted selection can drive alleles to extinction, regardless of their long-term benefit to a lineage. This is most readily seen in the case of a changing environment (Figure 2), where it has been noted in several contexts (Cvijović, et al. 2015, Gerland and Hwa 2009, King and Masel 2007, Masel, et al. 2007). Assume that a mutation arises in an environment in which it is beneficial and that the environment is constant for τ generations. Provided it survives genetic drift, the allele will increase in frequency following a logistic function and reach a frequency of one in approximately 2·ln(*Ns*)/*s* generations (Desai and Fisher 2007). Thus, if τ ≫ 2·ln(*Ns*)/*s*, then alleles will tend to arise and fix all in the same environment (Cvijović, et al. 2015). This provides a straightforward threshold, beyond which natural selection is blind to the allele’s long-term benefit. Of course, this threshold is derived under the assumption of a well-mixed population of constant size, and other factors such as demographic changes and population subdivision could substantially extend this upper bound. Still, these considerations demonstrate an inherent time-constraint imposed by evolution in finite populations, which only disappear as a mathematical artifact in infinite populations (Figure 2C).

**Figure 2.**
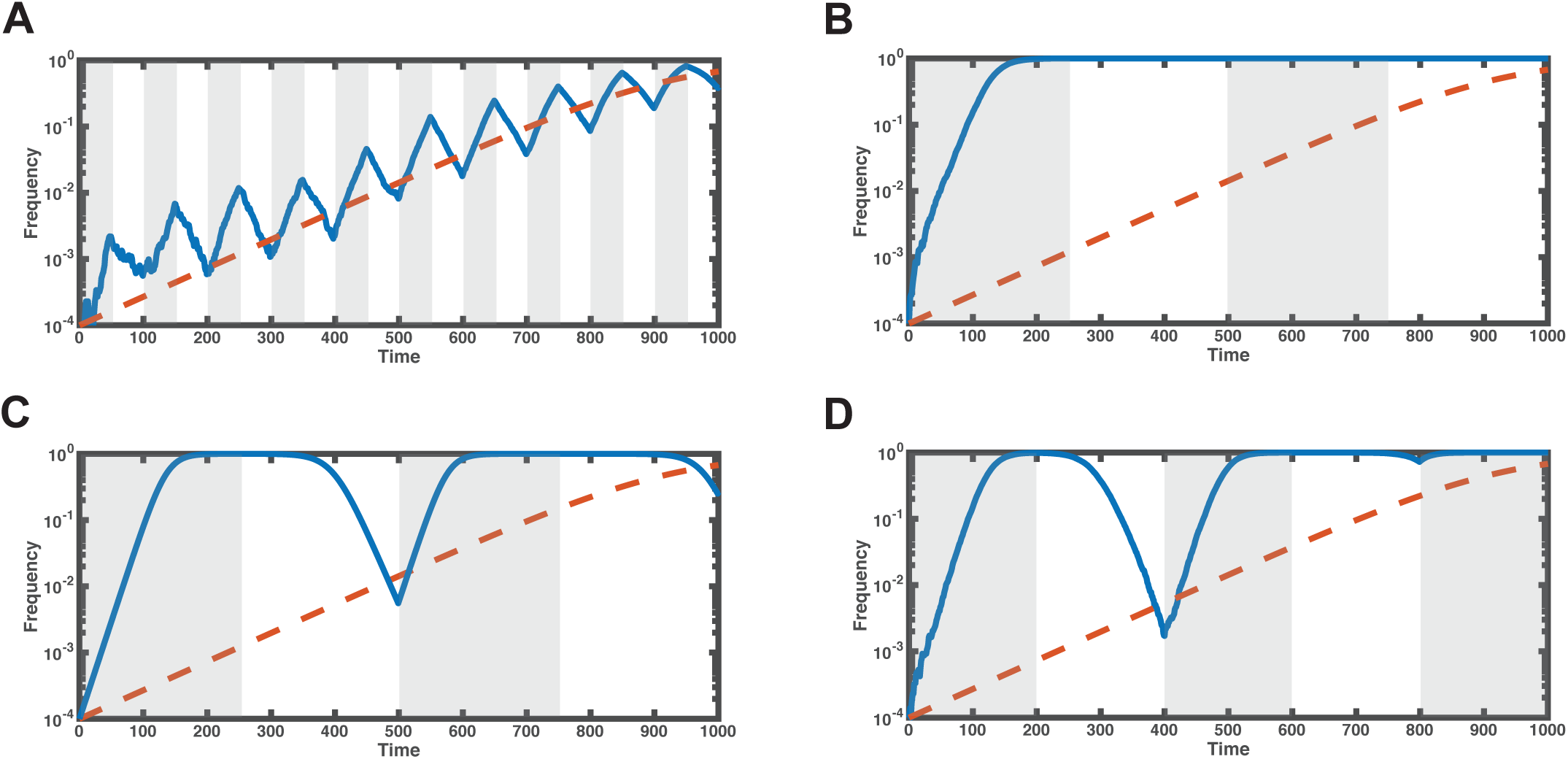
Limitations of fitness averages in finite populations. Evolution in a periodic environment results in four distinct regimes characterized by the relative timescale of natural selection (1/*s*) and environmental change (τ). Simulation results of the model described by Cvijović et al. (2015) are shown in blue and the expected change of an allele with the same average fitness effect in a constant environment is shown in red. Unless otherwise noted, simulations are conducted with an average selection coefficient of 0.05 and a population size of 100,000. Selection coefficients are held constant at ± 0.06 within each environment while the timescale of environmental change is varied (beneficial environmental epochs are shaded grey while deleterious epochs are unshaded). **A**. When the environment changes fast relative to changes in allele frequency (small sτ), the average change in allele frequency is well approximated by a fitness average like geometric mean fitness. **B**. When the environment changes slower than the time of fixation of an allele (large sτ), mutations tend to arise and fix all in the same environment, regardless of their average fitness effect. **C**. In infinite populations, averages like the geometric mean fitness are accurate regardless of the timescale of environmental change. This is an artifact of the fact that, in the absence of genetic drift, allele frequencies can become arbitrarily close to zero or one but never permanently achieve fixation or extinction. **D**. Average fitness breaks down when large fluctuations in allele frequency occur on a similar timescale to environmental change (intermediate sτ). This is due to the amplification of fluctuations by genetic drift whenever alleles reach very high or very low frequencies (Cvijović et al. 2015). Note that genetic drift occurring as the frequency of the allele approaches 1 causes it to respond only modestly to the second deleterious epoch. The allele subsequently achieved fixation much sooner than would be expected on the basis of its average fitness effect.

Similar limitations can be seen whenever the timescale of change in the fitness effects of an allele are greater than the time needed for natural selection to fix alleles conferring a short-term advantage. For example, models of multi-level selection become dominated by shortsighted selection of selfish phenotypes whenever group-level reproductive events are rare (Luo 2014). This breakdown in favor of shortsighted selection is analogous to that in variable environments (compare Figures 2B and 2D) and can be understood by considering the relative effects of individual and group selection on changes to allele frequency. Natural selection takes about s generations to double the frequency of a selfish trait within groups, where *s* denotes the within-group benefit of a selfish trait. On the other hand, increased rates of group reproduction in groups of non-selfish individuals will double the frequency of a cooperative trait after approximately *w·r* generations, where *r* is the group-level selection coefficient and *w* is the number of individual generations between group reproductive events. This heuristic reasoning implies that shortsighted selection in favor of a selfish trait will dominate allele frequency changes and preclude the evolution of cooperation whenever *s* ≫ *w·r*, which very closely matches results derived by formal analysis (Luo 2014).

### Beyond fitness averages

In addition to the role of extinction in tipping the outcome of selection toward shortsighted traits, studies explicitly modeling variability across a lineage have yielded a number of additional results that are not readily captured by Equation 1. Recently, Cvijović, et al. (2015) examined the case of a periodic environment that alternates between two states. An allele that is favored in one environment but disfavored in the next can follow unintuitive dynamics, particularly when large changes in allele frequency occur within environmental epochs. In the classic, lineage-invariant scenario discussed above, fixation of a neutral allele from a single starting copy requires traversing from a starting frequency of 1/*N* to a frequency of 1 by the action of genetic drift alone. In contrast, mutations in a fluctuating environment experience selective pressures continually, albeit of varying signs and intensities. This means that alleles can be driven to very high or very low frequencies by natural selection and then achieve fixation or loss due to genetic drift with far greater probability than predicted by Equation 1. This effect can cause the fixation probability of an allele to increase well beyond the neutral expectation of 1/*N*, even when alleles are neutral or deleterious on average. Furthermore, natural selection becomes less efficient at recognizing long-term fitness effects- causing mutations to behave as though they were effectively neutral, even when they are beneficial or deleterious on average. Finally, as populations become smaller or swings in frequency more dramatic, fixation can become independent of the average selection coefficient, creating conditions where the fixation probability is not even a monotonically increasing function of long-term fitness.

Another intriguing result emerges when the mean reproductive success across a lineage is held constant but its variance is altered. For example, Gillespie (1974) considered a model meant to capture spatial variation in the environment by relaxing the assumption of a Poisson-distributed number of offspring. Gillespie found that the natural way to quantify fitness is 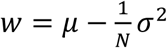 where μ is the mean number of offspring, σ^2^ is its variance, and *N* is the population size. A striking feature of this model is the appearance of population size in the definition of fitness, which suggests that the same allele can be favored or disfavored depending solely on the population size. This same sort of dependence on population size arises in a model of fluctuating environments (Takahata, et al. 1975), as well as in mutators (André and Godelle 2006, Raynes, et al. 2014, Wylie, et al. 2009). We emphasize that the population size dependence in the above examples is distinct from that of Equation 1, where population size influences the efficiency of natural selection but does affect its sign. Instead, variability in fitness across a lineage makes it possible that a subset of individuals will experience strong selective pressures that are not dominated by drift, even in small populations. This implies that genetic drift and natural selection do not, in general, scale according the relationship in Equation 1.

Perhaps the most intriguing feature of lineage variability is the possibility that the fate of an allele may not always be reducible to a selection coefficient at all. This is certainly the case for the evolution of mutation rate modifiers, where the succession of *de novo* beneficial and deleterious mutations results not only in variability in the distribution of offspring numbers across a lineage, but also in temporal autocorrelation in this distribution among the resulting sub-lineages (Figure 2C). Consequently, the offspring distribution is not only changing through time, but is also inherently linked to the underlying lineage dynamics. This implies that one is unable to define any selection coefficient for a mutator that predicts *P_fix._*, but must instead derive *P_fix_* directly under models that explicitly capture the dynamics of secondary mutations and clonal interference (Good and Desai 2016). Although one could then use *P_fix_* to retrospectively define an effective coefficient for any given population size using Equation 1 (Wylie, et al. 2009), it seems that one cannot generally define such a selection coefficient *a priori*. Moreover, even given such an effective selection coefficient, true *P_fix_* doesn’t scale with *N* in the manner predicted by Equation 1 (Good and Desai 2016). It remains to be seen whether a similar inability to reduce lineage fate to any effective selection coefficient might emerge in the context of variable environments and other examples of lineage-variable fitness effects.

## V. Implications for infectious disease evolution

One of the most promising applications of models considering lineage-variable fitness effects is in predicting and controlling the evolution of infectious diseases. Medically important traits such as pathogen virulence and drug resistance evolve rapidly and there has been considerable interest in the development of evolution-proof vaccines and antibiotics (Allen, et al. 2014, Day and Read 2016, Huijben, et al. 2013, Read, et al. 2011). Pathogen lineages experience a variety of extrinsic environmental changes including a dynamic immune response, a diverse set of tissues and hosts, and varying exposure to drugs. Additionally, since reproduction occurs both within and between hosts, multi-level selection can create conflicting selective pressures operating over different timescales (Kawashima, et al. 2009, Levin and Bull 1994). Finally, the dynamic immune response targeting antigenic epitopes has resulted in the selective pressures favoring mutator genes capable of immune evasion and antigenic evolvability (Deitsch, et al. 2009, Graves, et al. 2013, Moxon, et al. 1994). Variability across lineages therefore appears to be the rule rather than the exception in infectious disease evolution.

Predicting pathogen evolution and designing evolution-proof drugs will be greatly aided by models that combine the various selective pressures operating at different levels and timescales during the pathogen life-cycle. Traditional models have generally assumed that natural selection will favor traits that increase the long-term epidemiological success. For example, virulence is widely regarded as an adaptation to balance the increased rate of transmission by more aggressive diseases with the reduced duration of infection caused by host mortality or immune selection (Alizon, et al. 2009, Alizon and Michalakis 2015, Anderson and May 1982, Bull and Lauring 2014). However, the assumption that natural selection will maximize transmission success is analogous to selection maximizing other long-term measures of lineage success, like geometric mean fitness, and is therefore sensitive to the limitations discussed above (Figure 2). Specifically, shortsighted selection occurring within-hosts may act as a barrier for traits that could increase long-term transmission success (Levin and Bull 1994; Sidebar 2). Indeed, models that include mutation or competition between strains within-hosts or other ecological dynamics have demonstrated the inability of selection to maximize transmission success (Alizon, et al. 2013, Bonhoeffer and Nowak 1994, Day 2003).

There is broad support for the prediction that shortsighted selection and selection acting to increase traits that are beneficial on average can interact to shape infectious disease traits. For example, empirical studies in HIV (Alizon and Fraser 2013) and enteric bacteria (Giraud, et al. 2001) show how short-sighted selection can dominate patterns of evolution and lead to reductions in long-term transmission success. In *Salmonella enterica*, the need to maintain costly virulence factors that are susceptible to short-sighted selection for cheaters appears to have favored a strategy of cheater prevention that help to stabilize long-term infectivity (Diard, et al. 2013, Frank 2013, Mulder and Coombes 2013). Further theoretical progress on the role of interaction between the differing timescales of selection in pathogens could come from models that explicitly combine mechanistic within-host processes with long-term epidemiological dynamics (Coombs, et al. 2007, Day and Gandon 2007, Gilchrist and Coombs 2006, Mideo, et al. 2008). In addition, new experimental technologies such as lineage tracking of pathogens using barcode deep-sequencing (Blundell and Levy 2014, Levy, et al. 2015) offer exciting opportunities to measure selective pressures occurring within-hosts and integrate them with more traditional epidemiological data.

## VI. Conclusions

Despite historical emphasis on individual fitness effects shaping the fate of an allele, such a measure of fitness cannot always capture long-term evolutionary behavior when variability in fitness effects arise due to environmental, genetic or social interactions (Table 1, Figure 1). In some cases, averaging lineage-variable fitness across the various environmental, genetic, and social contexts an allele encounters allows for the application of classical population genetics results based on traditional notions of fitness related to individual survival and reproduction. However, this approach can fail in finite populations where an allele’s predicted fate can be interrupted by fixation or extinction due to shortsighted selection (Figure 2B). Furthermore, genetic drift and natural selection interact in unexpected ways when variability in fitness effects occurs over a comparable timescale to allele frequency (Cvijović et al. 2015, Figure 2D). More strikingly, examples from studies of mutation rate modifiers indicate that there may be no way to summarize the direction of natural selection on an allele without simply modeling its long-term lineage dynamics (Good and Desai 2016). Taken together, these findings may have particular relevance for the study of infectious pathogens, where alleles are likely to experience variability due to a combination of environmental, genetic, and social interactions.

Variability in the fitness effects of an allele challenge the conventional premise of population genetics that individual offspring number can be drawn from a fixed distribution for all members of a lineage (Figure 1). Cases where the typical assumption of a Poisson offspring distribution have been relaxed (Gillespie 1974) have yielded intriguing new evolutionary properties such as dependence on both the mean and variance in fitness effects and a critical effect of population size in determining whether an allele is beneficial. Other examples allow properties of the offspring distribution to vary in time, but still assume that the form of the distribution is fixed (Cvijović, et al. 2015). In yet other cases, it appears that allele frequency dynamics cannot always be reduced to one of independent draws from any offspring distribution, time-dependent or not. This effect is most recognizable in mutators, where the offspring distribution changes in a manner that is inseparable from the underlying lineage dynamics caused by secondary mutations and selection on sub-lineages (Figure 1C). Thus, while theoretical progress has been made in understanding processes where the offspring distribution takes on more general forms (Cannings 1974, Der, et al. 2011), we are still far from a population genetics theory with which to predict the fate of an allele in general scenarios of lineage-variable fitness effects.

Lineage variability also highlights the need for caution when interpreting the adaptive significance of biological traits in nature. Emphasis has often been placed on individual fitness effects at the expense of neglecting the ability of selection to favor traits that have longer term consequences on the fate of an allele (Williams 1966). Indeed, there are a plurality of definitions of fitness (Orr 2009) with each generalizing the concept of fitness under a particular source of lineage variability but none that appear sufficiently general to account for all examples. Caution is warranted when considering traits in the context of their long-term effects on a lineage, since such traits are inherently susceptible to shortsighted selection (Figure 2). Thus, while it is often safe to assume that selection will favor traits on the basis of extended fitness metrics, it is also important to consider the inherent limitations in the ability of natural selection to optimize any measure of fitness.

## Acknowledgements

The authors gratefully acknowledge Erol Akçay, Michael Desai, Steven A. Frank, Joanna Masel, Paul Rainey, Richard Watson, Eugene Raynes and other members of the Weinreich lab for numerous constructive comments on an earlier draft. C.J.G. is supported in part by NSF Graduate Research Fellowship 1644760, NSF 1501355 and NSF DGE 0966060. D.M.W. supported in part by NSF DEB-1556300 and NIH R01GM095728.

## Sidebar 1 – What is a lineage? (Typeset near ‘Introduction’)

We define a lineage as the full genealogy of descendent copies of an allele starting from the original copy and ending at its long-term fate: extinction, fixation, or maintenance as a stable polymorphism in the population. A traditional approach in population genetics has been to describe the long-term evolutionary fate of a mutant allele influencing some biological trait under the combined influence of evolutionary processes like mutation, genetic drift, migration, and natural selection. Using analytical approaches from stochastic process theory, this work seeks to calculate the probability that such an allele ultimately reaches a frequency of one, or achieves fixation, in the population. This approach places emphasis not solely on the individual reproductive process but also on the long-term fate of a genetic lineage carrying the mutation. It therefore captures a much larger class of phenomena where fitness may not be directly affected among individual carriers of an allele but the allele instead influences the statistical properties of a lineage (Wylie, et al. 2009). Our focus will be primarily on lineage of asexual haploid lineages, which are easier to analyze and depict. However, the approach and definition of a lineage given here extends naturally to sexual diploid organisms.

## Sidebar 2 – Geometric mean fitness *(Typeset near beginning of ‘Lineage-Variable Fitness Effects’)*

A widely appreciated result regarding adaptations to varying environments is the principle that natural selection will favor traits based on their geometric mean fitness. When reproductive success changes between generations, natural selection favors traits that increase the long-term geometric mean fitness (GMF). Reflecting the multiplicative nature of reproduction, GMF is the product of fitness in each generation, raised to the reciprocal of the number of generations. Algebraically, 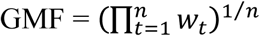, where *w_t_* is the Wrightian fitness of a trait in generation *t*. The same quantity can be expressed as a linear average over the natural log of this fitness value, 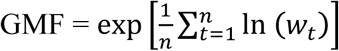 In practice, approximations are used such as 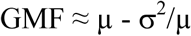, where μ is the arithmetic mean fitness and σ^2^ is the variance in fitness. This formula explicates the fact that natural selection favors increases in mean fitness, but also decreases in the variance of fitness. This implies that natural selection can be risk averse, favoring alleles with lower variance in fitness even at the expense of decreasing fitness on average.

## Sidebar 3 – Evolutionary game theory (Typeset near end of ‘Lineage-Variable Fitness Effects’or beginning of ‘Limitations of Fitness averages’)

Evolutionary game theory (Maynard Smith 1982) analyzes an interaction among a set of competing alleles or “strategies” and summarizes their effect in a matrix representing the fitness payoff of all pairwise competitions among competitors. Such a framework is most useful in the context of frequency dependent selection, where the fitness effects of an allele are not easily summarized by a constant selection coefficient. Such a framework provides a natural way to determine whether a new allele starting from a single copy will tend to increase in frequency or “invade” a population that is fixed for an alternative allele. This leads to the concept of an evolutionarily stable strategy or ESS, which is defined as a strategy or allele that cannot be invaded by any alternative strategy starting at an initially small frequency. The ability of an allele to invade, or invasion fitness, is a generalization of the notion of a selection coefficient to the case of frequency dependent selection and describes the long-term stability of a mutation against other competing mutations (Eshel, et al. 1998, Lehmann, et al. 2016). While there are notable exceptions (Traulsen and Hauert 2009, Traulsen and Nowak 2006), game theoretic models are typically deterministic and describe the central tendency for allele frequency change but not the statistical properties of lineages in finite populations.

**Glossary of terms** (*To appear adjacent to first use of each term or phrase*)

**Lineage-variable fitness effects:** Differing fitness effects of an allele between individuals due to genetic, social, or environmental interactions.

**Offspring distribution:** A discrete probability distribution that captures the stochasticity in an individual organism’s reproductive success.

**Cheater**: An individual that benefits from cooperative interactions of other individuals without itself contributing to the cost of cooperating.

**Frequency dependent selection:** A model in which the fitness of an allele depends on its frequency in the population as a consequence of interactions between organisms.

**Epistasis:** The phenotypic effect of a mutation varies with genetic context.

**Modifier loci:** Loci responsible for genetic properties of a genome, such as mutation rate, recombination rate and mutational robustness.

**Indirect selection:** Selection acting on a modifier locus mediated by genetic linkage with fitness effects at other loci in the genome.

**Clonal interference:** Competition between mutational independent asexual lineages, each carrying one or more beneficial mutations.

**Genetic drift:** Stochastic variation in allele frequency as a consequence of stochasticity in reproduction inherent in finite populations.

